# Benefits And Limitations Of Three-Dimensional Printing Technology For Ecological Research

**DOI:** 10.1101/283895

**Authors:** Jocelyn E. Behm, Brenna R. Waite, S. Tonia Hsieh, Matthew R. Helmus

**Affiliations:** Integrative Ecology Lab, Center for Biodiversity, Department of Biology, Temple University, Philadelphia, PA, USA; Department of Ecological Science – Animal Ecology, VU University Amsterdam, Amsterdam, the Netherlands; Department of Biology, Temple University, Philadelphia, PA, USA

**Keywords:** 3D models, additive manufacturing, *Anolis sagrei*, clay model, Curaçao, Maya Autodesk, sustainability

## Abstract

**Background:** Ecological research often involves sampling and manipulating non-model organisms that reside in heterogeneous environments. As such, ecologists often adapt techniques and ideas from industry and other scientific fields to design and build equipment, tools, and experimental contraptions custom-made for the ecological systems under study. Three-dimensional (3D) printing provides a way to rapidly produce identical and novel objects that could be used in ecological studies, yet ecologists have been slow to adopt this new technology. Here, we provide ecologists with an introduction to 3D printing.

**Results:** First, we give an overview of the ecological research areas in which 3D printing is predicted to be the most impactful and review current studies that have already used 3D printed objects. We then outline a methodological workflow for integrating 3D printing into an ecological research program and give a detailed example of a successful implementation of our 3D printing workflow for 3D printed models of the brown anole, *Anolis sagrei,* for a field predation study. After testing two print media in the field, we show that the models printed from the less expensive and more sustainable material (blend of 70% plastic and 30% recycled wood fiber) were just as durable and had equal predator attack rates as the more expensive material (100% virgin plastic).

**Conclusions:** Overall, 3D printing can provide time and cost savings to ecologists, and with recent advances in less toxic, biodegradable, and recyclable print materials, ecologists can choose to minimize social and environmental impacts associated with 3D printing. The main hurdles for implementing 3D printing – availability of resources like printers, scanners, and software, as well as reaching proficiency in using 3D image software – may be easier to overcome at institutions with digital imaging centers run by knowledgeable staff. As with any new technology, the benefits of 3D printing are specific to a particular project, and ecologists must consider the investments of developing usable 3D materials for research versus other methods of generating those materials.

## Background

Ecologists exhibit exceptional creativity and ingenuity in designing new tools and equipment for their studies, often incorporating and repurposing technology from other fields. For example, unique solutions have been devised for tracking animals (backpack-mounted radio transmitters [1]), tracking seeds (fluorescent pigments [2]; seed tags [3]), catching animals (pit-less pitfall traps [4]), containing or restraining difficult-to-hold specimens (squeeze box for venomous snakes [5], ovagram for amphibian eggs [6]), and remotely collecting data or samples (frog logger [7]; hair trap [8]), among countless others. Because many ecological studies require customized equipment, ecologists are no strangers to building the contraptions necessary for conducting their research, and the weeks leading up to and during field seasons and lab experiments often involve multiple trips to hardware stores and craft shops.

Despite the high level of creativity and adaptability exhibited by ecologists, there is one technology that ecologists have been slower to adopt relative to other fields: three-dimensional (3D) printing. Additive layer manufacturing, or 3D printing, is the layering of material by a computer-controlled machine tool to create an object from a digital file that defines its geometry [9]. Most objects are printed in plastic, but newer print materials such as metal, wood, or other composites are increasingly common in consumer applications. In the recent past (i.e., before 2010), 3D printing was cost-prohibitive and limited in availability, but it is now affordable and accessible to budget-conscious ecologists. Many research institutions have at least one 3D printing center and 3D printing services are available to all online. Other fields, such as the health sciences, have readily adopted 3D printing into their research [e.g., 10], but it is as of yet an untapped technology that ecologists can exploit to their advantage.

Recent studies have highlighted the benefits of 3D printing in terms of cost and time efficiency [11, 12], yet ecologists wanting to implement 3D printing for the first time must still traverse a steep learning curve. Our goal here is to flatten the curve and provide ecologists with a general but sufficient background in 3D printing technology to know what considerations are important when approaching a 3D printing project. In this article, we provide an overview of how 3D printing has been adopted by fields related to ecology. We highlight areas of ecological research where we believe 3D printing has the promise to be most effective, and provide a methodological workflow for integrating 3D printing into ecological studies. We illustrate this workflow using an example from our own work, which includes the obstacles we encountered and the solutions we devised. Finally, we conclude with important environmental sustainability considerations.

### Overview of 3D printing in fields related to ecology

Two disciplines that were early adopters of 3D printing technology and have strong connections to ecology are biomechanics and natural history curation. Below we provide examples of 3D printing implementations in these fields to provide ecologists with ideas of what is possible.

The aim of biomechanics is to understand the movement and structure of living organisms integrating across physics, engineering, physiology, and ecology. In biomechanics, 3D printing is used to test how the shapes of particular appendages or biological structures function in the physical environment without having to use live organisms. For example, 3D printed models of the sand-burrowing sandfish lizard’s (*Scincus scincus*) respiratory system made it possible to study why it does not inhale sand in ways that are impossible with an actual respiratory system [13]. In studies of fluid dynamics, 3D printed models of swift (*Apus apus*) wings and bodies of echolocating bat species permitted tests in water and wind tunnels respectively to understand how morphology influences species’ movements [14, 15]. In other applications, biomechanical theory is tested by attaching 3D printed structures to robots. In a study of underwater burrowing mimetics in bivalves, Germann *et al.* [16] used mathematical models to design a bivalve shell which was 3D printed and incorporated into a burrowing robot. In other studies, evolutionary optimization models are used to design the shape of anatomical structures. Then, 3D prints of the modeled and naturally occurring structures are compared in performance tests to understand the evolutionary limitations species face in structural adaptation [17, 18]. For these studies, 3D models enabled scientific inquiry, as manipulating live animals would have been challenging to impossible.

In the field of natural history curation, 3D printing increases the speed at which discoveries are made, and the rate at which data and resources are shared across natural history collections [19]. In paleontology, the reconstruction of complete skeletons is often impaired by the recovery of incomplete remains at dig sites. Mitsopoulou *et al.* [20] used mathematical allometric scaling models to calculate the dimensions of bones missing from the remains of a dwarf elephant (*Paleoloxodon tiliensis*) recovered from Charkadio Cave on Tilos Island, Greece. From these analyses, a 3D model was printed to allow the complete skeleton to be assembled. In addition, 3D technology also facilitates the sharing of museum material without having to loan valuable specimens, making it possible to construct complete skeletons using partial skeletons from multiple separate collections [21]. In fact, museums have been quick to adopt 3D technology because it vastly improves the rate at which collections are shared. The exchange of 3D-printed specimens facilitates crowd sourcing for specimen identification; access to high-quality replicas of endangered, extinct, or otherwise valuable and/or fragile specimens; and printed specimens can even be used in a field setting for species identification [21, 22]. Museums are increasingly accepting deposits of 3D printed material for rare and/or difficult to access specimens. Lak *et al.* [23] employed 3D technology to describe two new damselfly species that were preserved in amber. Because it is difficult to physically extract amber-encased specimens without damaging them, the team used phase contrast X-ray synchrotron microradiography to make 3D images of the specimens, and deposited the 3D prints in several museums. Finally, 3D technology also accelerates the flow of information for education and outreach. For example, Bokor *et al*. [24], developed a classroom exercise where students print fossilized horse teeth and examine how the teeth changed over time with respect to changing climate.

### Integration of 3D printing in Ecology

While ecologists have used 3D printing in a variety of applications (Table 1), there are four areas where we view 3D printing to be the most impactful: behavioral ecology, thermal ecology, building customized equipment, and enhancing collaboration.

**Table 1:**
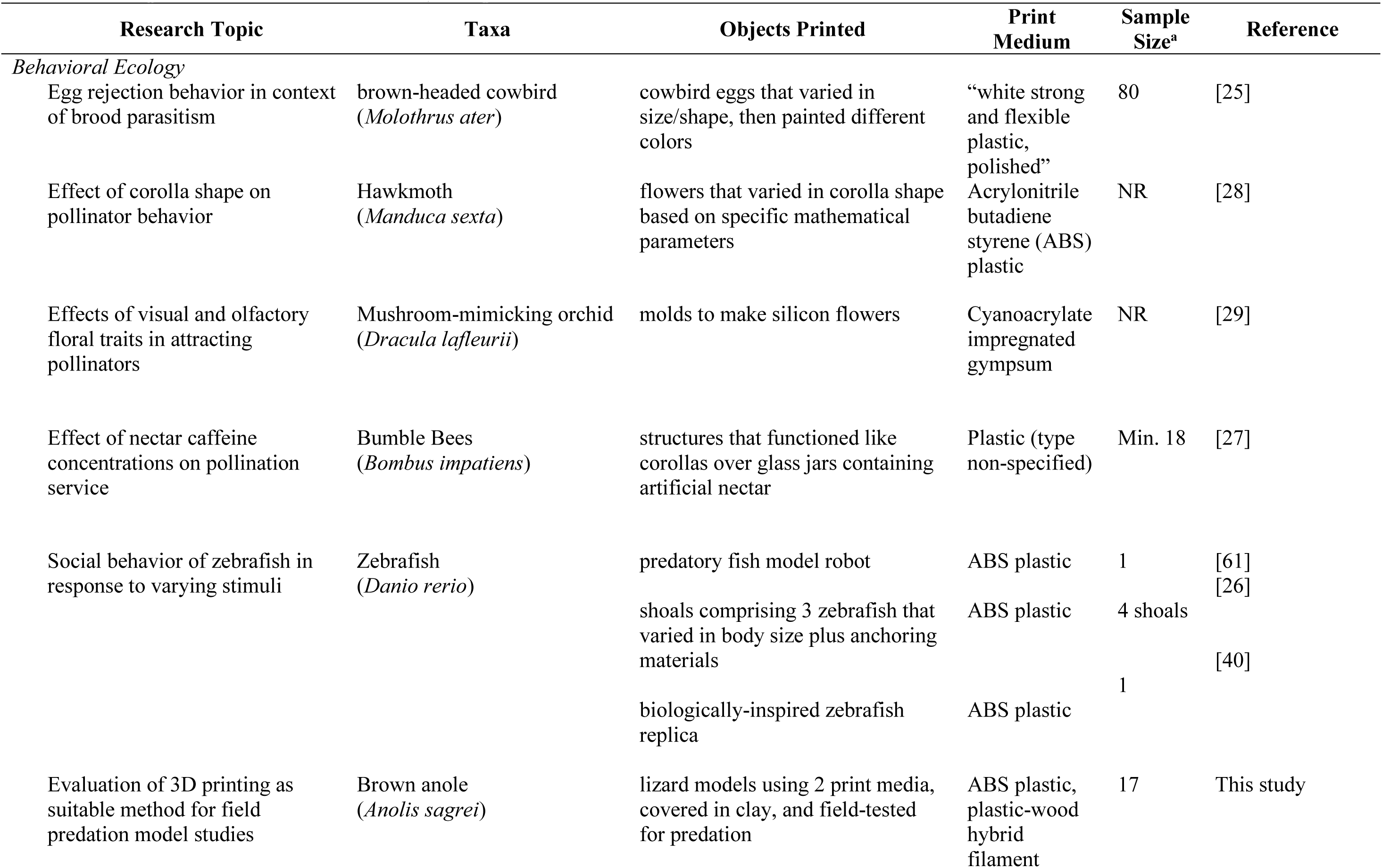
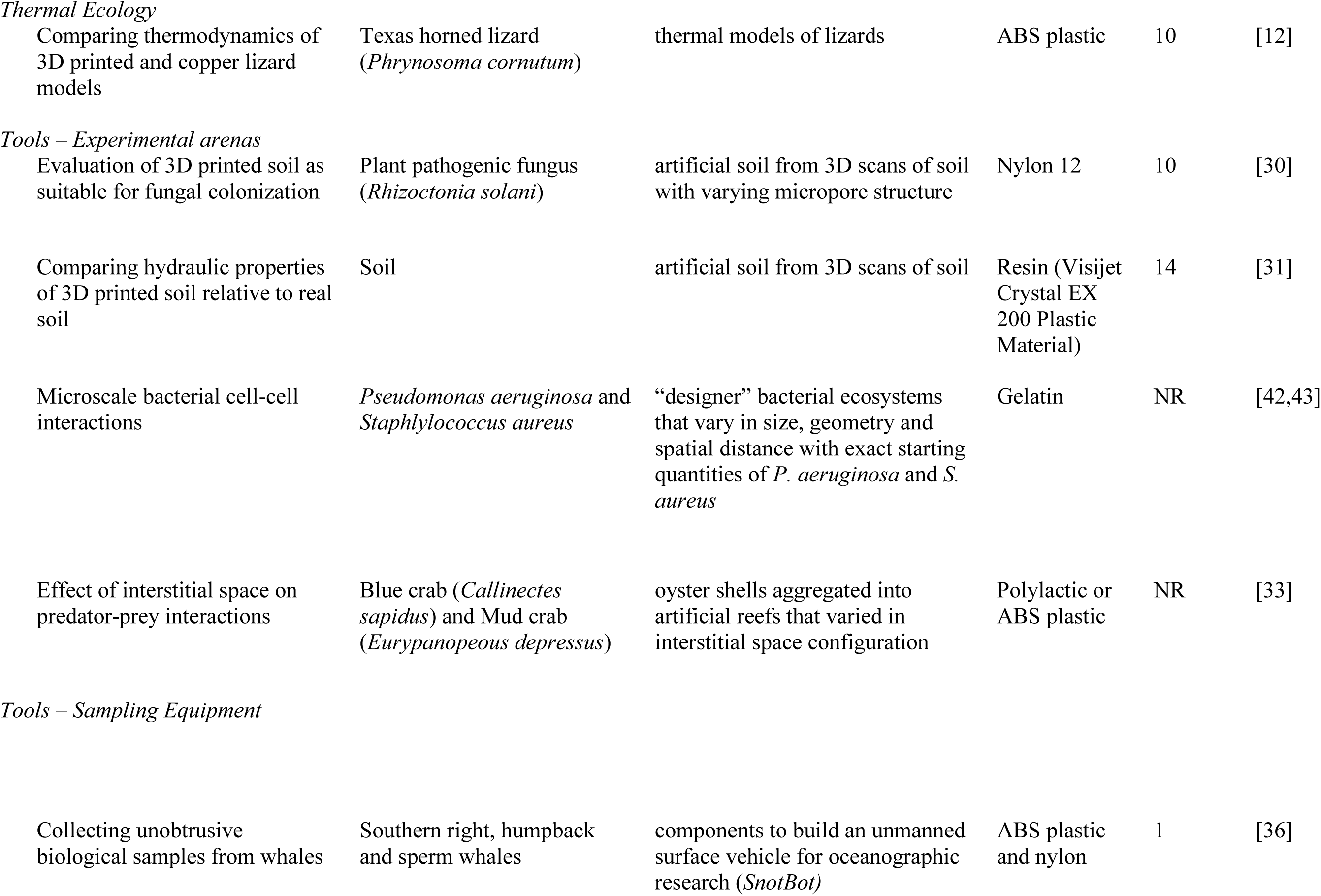
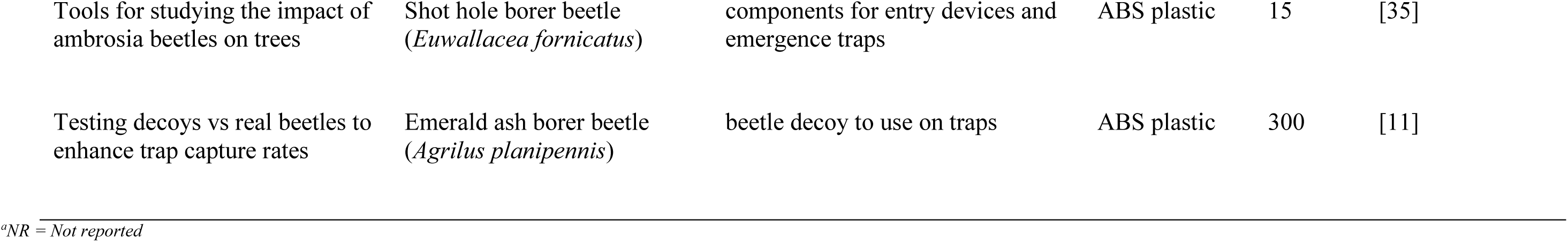
Ecological studies that have used 3D printing

| Research Topic | Taxa | Objects Printed | Print Medium | Sample Size<sup>a</sup> | Reference |
| --- | --- | --- | --- | --- | --- |
| *Behavioral Ecology* |  |  |  |  |  |
| Egg rejection behavior in context of brood parasitism | brown-headed cowbird (*Molothrus ater*) | cowbird eggs that varied in size/shape, then painted different colors | “white strong and flexible plastic, polished” | 80 | [25] |
| Effect of corolla shape on pollinator behavior | Hawkmoth (*Manduca sexta*) | flowers that varied in corolla shape based on specific mathematical parameters | Acrylonitrile butadiene styrene (ABS) plastic | NR | [28] |
| Effects of visual and olfactory floral traits in attracting pollinators | Mushroom-mimicking orchid (*Dracula lafleurii*) | molds to make silicon flowers | Cyanoacrylate impregnated gypsum | NR | [29] |
| Effect of nectar caffeine concentrations on pollination service | Bumble Bees (*Bombus impatiens*) | structures that functioned like corollas over glass jars containing artificial nectar | Plastic (type non-specified) | Min. 18 | [27] |
| Social behavior of zebrafish in response to varying stimuli | Zebrafish (*Danio rerio*) | predatory fish model robot | ABS plastic | 1 | [61] |
|  |  | shoals comprising 3 zebrafish that varied in body size plus anchoring materials | ABS plastic | 4 shoals | [26] |
|  |  | biologically-inspired zebrafish replica | ABS plastic | 1 | [40] |
| Evaluation of 3D printing as suitable method for field predation model studies | Brown anole (*Anolis sagrei*) | lizard models using 2 print media, covered in clay, and field-tested for predation | ABS plastic, plastic-wood hybrid filament | 17 | This study |
| *Thermal Ecology* |  |  |  |  |  |
| Comparing thermodynamics of 3D printed and copper lizard models | Texas horned lizard (*Phrynosoma cornutum*) | thermal models of lizards | ABS plastic | 10 | [12] |
| *Tools – Experimental arenas* |  |  |  |  |  |
| Evaluation of 3D printed soil as suitable for fungal colonization | Plant pathogenic fungus (*Rhizoctonia solani*) | artificial soil from 3D scans of soil with varying micropore structure | Nylon 12 | 10 | [30] |
| Comparing hydraulic properties of 3D printed soil relative to real soil | Soil | artificial soil from 3D scans of soil | Resin (Visijet Crystal EX 200 Plastic Material) | 14 | [31] |
| Microscale bacterial cell-cell interactions | *Pseudomonas aeruginosa* and *Staphylococcus aureus* | “designer” bacterial ecosystems that vary in size, geometry and spatial distance with exact starting quantities of *P. aeruginosa* and *S. aureus* | Gelatin | NR | [42,43] |
| Effect of interstitial space on predator-prey interactions | Blue crab (*Callinectes sapidus*) and Mud crab (*Eurypanopeus depressus*) | oyster shells aggregated into artificial reefs that varied in interstitial space configuration | Polylactic or ABS plastic | NR | [33] |
| *Tools – Sampling Equipment* |  |  |  |  |  |
| Collecting unobtrusive biological samples from whales | Southern right, humpback and sperm whales | components to build an unmanned surface vehicle for oceanographic research (*SnotBot*) | ABS plastic and nylon | 1 | [36] |
| Tools for studying the impact of ambrosia beetles on trees | Shot hole borer beetle (*Euwallacea fornicatus*) | components for entry devices and emergence traps | ABS plastic | 15 | [35] |
| Testing decoys vs real beetles to enhance trap capture rates | Emerald ash borer beetle (*Agrilus planipennis*) | beetle decoy to use on traps | ABS plastic | 300 | [11] |

<sup>a</sup>NR = *Not reported*

The main goal of behavioral ecology is to understand how ecological and evolutionary forces shape behavior. In addition to observational studies, behavioral ecology research can involve manipulations of environmental conditions to test hypotheses. For testing hypotheses in both lab and field conditions, 3D printing may be incredibly useful for making precise, repeatable models. Three-dimensional printing has already been used to create precise models of bird eggs to test egg rejection behavior in the context of brood parasitism [25], zebrafish shoals to test the effect of body size on zebrafish shoaling preferences [26], and artificial flower corollas to test the effect of floral traits on pollinator visitation [27–29] (Table 1). In these studies, 3D printing was chosen for its ability to create identical experimental stimuli because alternative methods, such as constructing models by hand, could introduce unintentional variation that makes it difficult to determine whether study subjects are responding to intentional or unintentional variation in experimental stimuli. In addition, 3D printing is often a faster method for creating models than making them by alternative methods [12]. There may be scenarios where 3D printing will not produce more biologically accurate models than other methods, but in many cases, 3D printing will increase the types of behavioral questions that can be asked [25]. Within the field of behavioral ecology research, 3D printing can be used to test myriad behaviors including predation (see Results), reproduction, foraging, social interactions, and defense in both aquatic and terrestrial habitats.

Thermal ecology is focused on understanding how organisms are influenced by the temperature profile of their environment. A major challenge of thermal ecology research is constructing models that accurately replicate the thermal properties of a study organism. Copper models are often used, however, recent work demonstrated that 3D printed plastic models were cheaper and faster to construct and exhibited no difference in thermal properties compared to standard copper models (Table 1) [12]. This, as well as the need for high numbers of identical models, suggests that 3D printed models may make thermal ecology research more accessible.

Perhaps 3D printing will be the most helpful to the widest number of ecologists because it provides a method for constructing customized equipment such as tools and experimental habitats or mesocosms. In the field of soil ecology, 3D printing has been used to print artificial soil structures which accurately replicate the macropore structure of soil (Table 1) [30, 31]. These artificial soils are ideal replicate experimental mesocosms for soil macro- and/or microorganisms. Structures designed for other studies could be repurposed by ecologists as experimental habitats such as artificial gravel beds originally designed for testing water flow patterns [32] and artificial oyster shell reefs used to test how habitat complexity influences predation rates [33].

Opportunities for printing tools are limited primarily by the ecologists’ imagination and range from simple structures to complex moving machines [34]. On the low-complexity end of the spectrum, 3D printing has been used to sample two difficult-to-catch, invasive, tree-boring beetle species that cause significant damage. Three-dimensional printed emergence traps make it possible to effectively trap and census invasive ambrosia beetles *(Euwallacea fornicates*) as they emerge from trees [35], while 3D printed decoys placed on standard beetle traps enhanced capture rates of invasive emerald ash borer beetles (*Agrilus planipennis*) [11]. In a more complex application, whale researchers used 3D printing to build an unmanned surface vehicle named *SnotBot* which allows scientists to get close enough to whales to collect biological samples (Table 1) [36]. There are ample opportunities for ecologists to design tools to aid in data collection, sample processing, organism containment, and even organization of field or lab spaces.

From the examples provided above, designing custom materials certainly benefits scientists within the context of a particular study. However, the use of 3D technology also provides a mechanism for collaboration that extends beyond the limits of a single study. Ecological studies that are replicated across systems, geographic boundaries, latitudinal gradients, etc., are a powerful method for testing ecological theory [37]. The use of 3D technology facilitates these broad-scale studies through the sharing of identical tools, models, and/or equipment that can be used in multiple systems. For example, 3D printed models of brown-headed cowbird (*Molothrus ater*) eggs [25] and Texas horned lizards (*Phrynosoma cornutum*) [12] can be used to test patterns of brood parasitism and thermal tolerances, respectively, across their geographic ranges. Similarly, for widespread invasive species like the emerald ash borer, sharing effective trap methodology [11] among scientists and agencies can potentially accelerate the rate at which the impact of the species is mitigated. In addition, 3D technology provides a useful platform for ecologists who would like to incorporate citizen scientists into a research program. Indeed, effective sampling technologies that can be disseminated electronically are ideal for citizen science, and increase the speed at which consistent data can be collected [38].

## Methods

Below we describe a general workflow to use when embarking on incorporating 3D printing into ecological research. Essentially, once an ecologist has identified the object to be printed, the 3D printing process involves creating a printable 3D digital image file of the object, selecting an appropriate print media, and then printing draft and final versions of the object (Figure 1). To be clear, details specific to each project and available resources will need to be explored and fine-tuned along the way. However, our workflow highlights the major steps and aspects to consider at the onset.

**Fig. 1.**
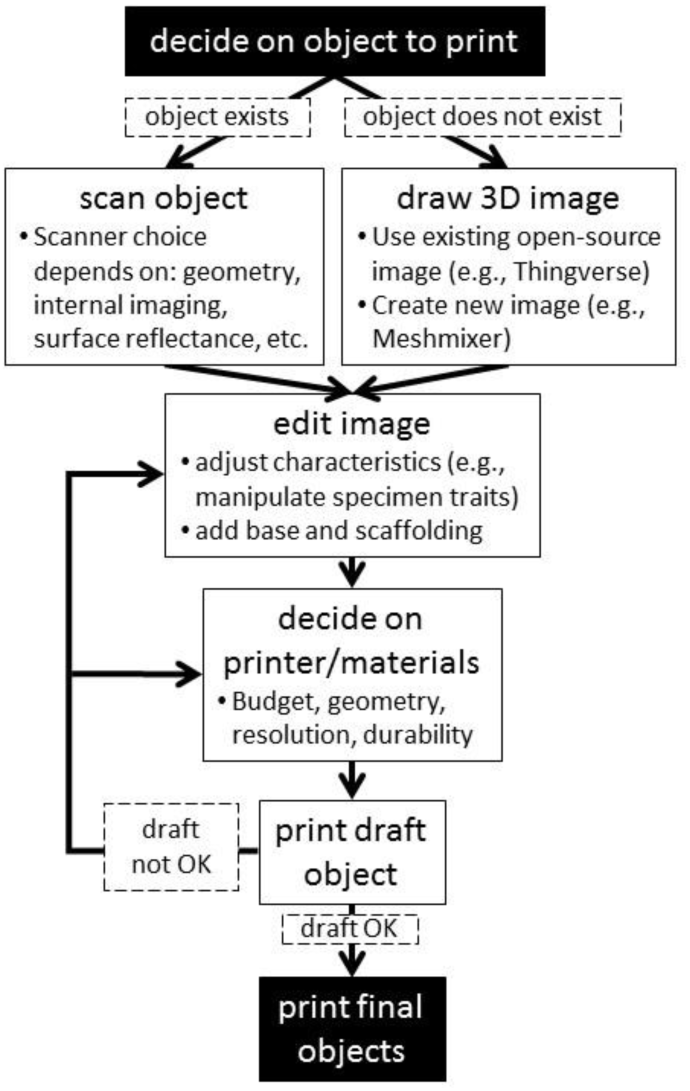
Workflow illustrating steps for integrating 3D printing in ecological research

### Make a digital object file

The first step is to generate a digital file of the object to be printed, which can be accomplished by creating a digital file of the image from scratch, converting a 2D image (e.g., photograph) into a 3D image, scanning an existing 3D object, or using an existing 3D file. All digital 3D files require use of software specifically for editing 3D images (Additional File 1). The most common 3D image file format is an STL file and is used by many software packages. Depending on the image generating methods used and the types of modifications needed, there may be a significant learning curve to attain the necessary level of proficiency on the software. This is especially true for creating a 3D image completely from scratch (see below). In our experience, however, we scanned an existing object and an undergraduate student was able to work together with the printing center staff to learn the software and manipulate the image within two months.

Before trying to create the image from scratch or scan an existing image, it may be worthwhile first to check the many libraries of 3D imagery that are available online (Additional File 2). It is possible that a digital 3D file of a similar object has already been created and can be downloaded potentially for free, ready to be printed. Even if the file in an online library is not exactly perfect, it can be manipulated using 3D software (Additional File 1), which, depending on the modifications needed, may be a more efficient use of time than scanning an image or trying to draft an image from scratch.

If a suitable digital 3D file is not available, but the object to be printed is in the ecologist’s possession, it is possible to use a 3D scanner to make a digital 3D image of the object, similar to how a flatbed scanner makes a digital 2D image of an object. There are various types of scanners, and it is necessary to choose a scanner that can accurately capture the level of detail from the object being scanned needed for the project. Laser scanners, structured light scanners, and even smart phone apps, can be used to create lower resolution scans of an object’s external features. Laser scanners were used to scan Texas horned lizards that were frozen in realistic positions for a thermal ecology study (Makerbot Digitizer 3D, Makerbot, New York, USA) [12], and oyster shells for a biomechanical predation study (Vivid 9i, Konika Minolta Inc., Tokyo, Japan) [33]. For more complex and fine scale objects with both internal and external features like soil micropore structure, a method like X-ray microtomography is more appropriate (HMX 225, Nikon Corp., Tokyo, Japan) [30].

If the object to be printed is not in the ecologist’s possession, it is possible to design the object using 3D drafting software (Additional File 1), with the time investment being proportional to the researcher’s proficiency on the software and the complexity of the object. Using photogrammetry, photos can be digitized and 2D x,y coordinates from the photo converted into a 3D image [25, 39]. Photogrammetry may be the easiest and most cost effective method, especially if a scanner is not available. Alternatively, mathematical formulae may be used to generate different shapes, such as the surface of a bird egg [25] or the curvature of a flower corolla [28]. Finally, it is possible to draft the object completely from scratch [e.g., 35], although a higher proficiency on the appropriate drafting software is necessary (Additional File 2).

Once a digital 3D image file is in hand, it will likely need to be edited and customized for the particular study. For example, in the brood parasitism study, the 3D image of the bird egg was edited to make it hollow so that the printed versions could be filled with water so their weight and thermal properties more closely matched a real bird egg [25]. Similarly, in the thermal ecology study, the 3D image of the Texas horned lizard was edited to include a well in the underside that fit a small environmental sensor (iButton) for measuring temperature [12]. Object size can also be manipulated and various polygons added to include additional structures.

Depending on the type of printer and material used, the image may need to be edited to make printing possible and to efficiently use printing material. Non-manifold geometry errors (i.e., geometry that cannot exist in the real world) can be common in scans made on biological objects and must be corrected in order avoid fatal printing errors. Most 3D file manipulation software allows for these corrections (Additional File 1). Because most printers print the object from the bottom up layer-by-layer, any appendages or protrusions that extend out much wider than the bottom layer may need added scaffolding to make the print possible. This scaffolding is removed after printing is completed with varying degrees of effort depending on the design and print material. In addition, if the object is not flat, it will likely need a flat base added to make it printable. If multiple copies of the object are to be printed, it may be possible to rotate or stack them so that several copies can be printed simultaneously. This method ensures efficient use of printing platform space and materials.

### Printer and printing material

There is a wide range of 3D printers that use various printing technologies and materials, and a comprehensive review of all printer types is beyond the scope of this article. Here, we focus on the printers and materials likely to be most useful to ecologists. Many factors must be weighed when choosing a printer and printing material for a project, such as cost, material durability, printed surface quality, timeframe for printing, and color. The most ubiquitous printers that are common on university campuses and also through commercial online printing services typically use either plastic-based filament or resin as the print material. Filament is hard plastic stored on spools that is melted and deposited as beads or streams during printing that quickly re-harden into layers to form the object. Resin is a polymer liquid that is layered and solidified with UV light. Both come in a range of colors; filament is often cheaper but leads to a lower resolution print with printed bands more prominent on the finished object, however if needed there may be applicable surface finishing methods for smoothing out these bands, like using acetone vapor. Filament may also be less durable for some applications and cracks can form between layers if the object is subjected to physical stress. Finished resin products are generally smoother, can be printed at higher resolution, are more durable, and have the surface quality of a store-bought plastic item.

Both filament and resin have been used for printing low and high resolution ecological models, respectively. For example, acrylonitrile butadiene styrene (ABS), a type of filament, was used for printing artificial flowers [28], artificial zebrafish [26, 40], and models of lizards [12], while resin was used for printing artificial soils with fine-scale pore structure in a hydrology study [31]. It is also worth considering the type of scaffolding involved with a specific printer/ print material combination. For some printing set-ups, the scaffolding is the same material as the printed object, which means the scaffolding must be physically cut off, creating opportunities to damage the printed object. Other printers are capable of dual or multi-extrusion, meaning they can print using different materials simultaneously. In this case, the scaffold material differs from the print material and can be dissolved after printing in a chemical solvent solution.

More high-tech printers capable of printing even finer-scale and more-detailed objects use a powder based print material which is converted into a solid plastic with a laser. An advantage of this print material is that little scaffolding is needed and extra powder can quickly be removed by shaking or brushing. This media was used to print soil pore microstructure at the scale of micrometers [30]. These artificial soils were printed using Nylon 12, a material that can be autoclaved, which makes it possible to reuse the soils for multiple experiments [30]. Although most standard printing materials are various types of plastic, there are a handful of products that include other materials like wood, rubber, and metal. At least one biodegradable plastic filament also exists: a polylactic acid (PLA) made from corn starch [34, 41]. Perhaps the most high-tech printers that have been used for an ecological application printed gelatin-based designer bacterial ecosystems that varied in geometry and spatial structure in order to study cell-to-cell interactions [42,43; Table 1], but this technology is not readily available to most ecologists.

### Printing

Once the 3D image has been drafted and edited, and the printer and print materials have been selected, a test round of printing is necessary before moving to the final round. Printing a test object makes it possible to identify errors with the 3D image file, compare print materials and confirm the material choice, and gain an estimate of the amount of time required for printing *en masse*. After all aspects of the printing project have been approved, the final prints can proceed. Following printing, various post-processing stages will likely need to occur, such as removing scaffolding, painting, adding clay, and/or assembling pieces.

## Results

Here we provide an example of a successful attempt to integrate 3D printing into an ecological project following the workflow outlined above. We include the obstacles encountered along the way as a useful case study for other ecologists. Note, we used equipment (scanners and printers) and expertise from two (out of the four) 3D printing centers at our institution. For ecologists with fewer onsite resources, online resources and resources at collaborating institutions may be useful.

Clay animal models have long been used in ecological field research to infer predation rates by free-ranging predators on prey. In this methodology, animal models are constructed from plasticine modeling clay and then placed in the field for a fixed time period. Because the clay does not harden, predation attempts leave marks in the clay, making it possible to score models for evidence of predation. Early work used this method to study how body coloration affected predation rates in snakes [44, 45]. Since then, clay models have been used in predation studies to represent a wide range of taxa including frogs [46], salamanders [47], lizards [48], and insect larvae [49].

In many of these studies, models are constructed by hand either completely or nearly completely from clay [e.g., 45,47,49,50]. In other studies, silicon molds are made from preserved specimens, which are then used to make models either directly out of clay [51], or out of plaster which is then covered with clay [52]. These methods clearly produce models that elicit responses in predators, however, producing the models in this manner can be time consuming as studies may use upwards of 100 models. In addition, modifying the models in a precise manner to test the effects of prey traits on predation is difficult. The repeatability, speed, and precision of 3D printing make it highly applicable to field studies of predation using models. We first explored the ease of creating a 3D scan of a preserved lizard specimen, and then used software to modify its body size. We then tested the durability of two print materials and two model sizes in a field predation study.

### Making the lizard model

We used two methods, a structured light scanner (David SLS-2 3D Scanner, HP Inc., Palo Alto, CA, USA) and a laser scanner (NextEngine2020, NextEngine, Inc., Santa Monica, CA, USA), to make 3D scans of a preserved male *Anolis sagrei* lizard. Structured light scanners operate by projecting light patterns onto the object being scanned and analyzing the pattern’s deformation with a camera. The laser scanner we used boasts new technology consisting of more sophisticated algorithms and multiple lasers which scan in parallel, yielding more data points and an overall more accurate scan. Both scanners are designed to scan 3D objects, but because they use different technologies to do so, one scanner may be more effective for scanning a particular object. Regardless of the number of scans or angle of rotation, the structured light scanner’s software was not able to converge the multiple scans into a single image of our anole, likely due to the complexity and high reflectance of the preserved specimen’s skin. The laser scanner, however, was able to produce a digital 3D image of the specimen within about 90 minutes, and we used this file going forward. The laser scanner was most successful when the lizard specimen was positioned in a vertical rather than flat manner using an Extra Part Gripper (NextEngine, Inc., Santa Monica, CA, USA; Figure 2A).

**Fig. 2.**
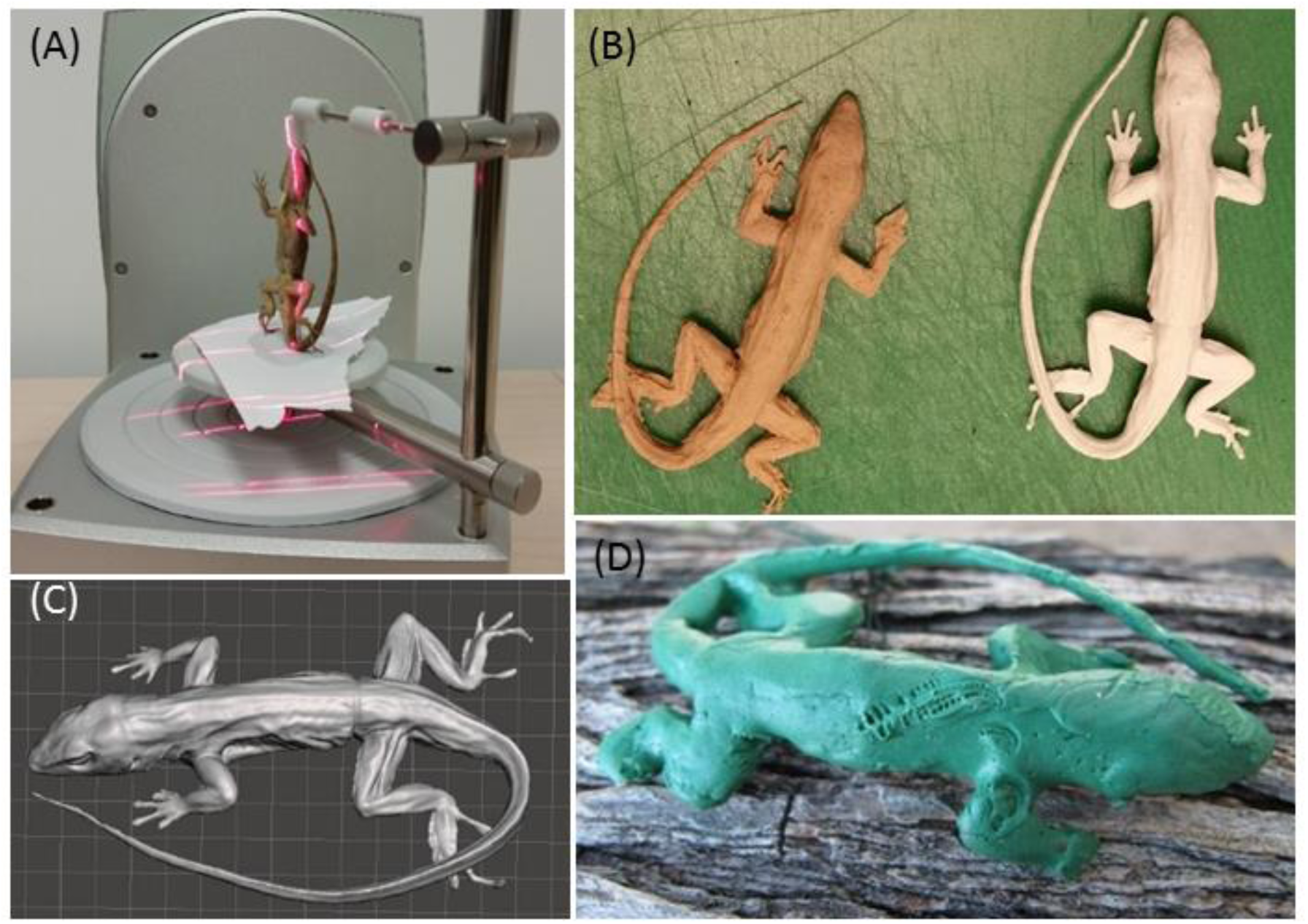
Construction of a 3D printed lizard predation model a) Successful laser scanning setup of preserved brown anole (*Anolis sagrei)* specimen in vertical orientation; b) 3D image of scanned anole viewed in Meshmixer software and later edited in Maya; c) 3D printed plastic-wood hybrid (left) and ABS plastic (right) anole models; d) clay covered model on a branch in the field with bite marks likely from a lizard predator (*Cnemidophorus murinus murinus*).

We used Maya software (Autodesk, San Rafael, CA, USA; Additional File 2) to edit the scanned image (Figure 2B) of the lizard specimen to attain three goals. First, to make the lizard scan possible to print, we had to edit the non-manifold geometry errors that arose due to the scanning process. Second, we manipulated the size of the lizard to test whether different printing materials were durable for both large and small prints. The large lizard was 25% larger than the original (snout vent length = 60 mm). Finally, we added a hollow horseshoe-shaped tube in the ventral side of the body cavity for looping a small wire through in order to anchor the models to branches in the field. The final file we used to print the lizards is available from the authors.

### Print material and printing

We tested two types of filament print media: plastic (ABS-P430 plastic in ivory, Stratasys, Eden Prairie, MN, USA) and plastic-wood hybrid (Woodfill by ColorFabb, Belfeld, the Netherlands). ABS exceeded the Woodfill in cost and perceived durability, yet Woodfill was a more sustainable option as it is made of 30% recycled wood fibers. During our test print stage, we learned we needed to add a base to our digital 3D image file for the Woodfill prints because the scanned image was not flat which made it difficult to print. We did not need to edit it for the ABS print because the scaffold base dissolved.

After we finalized our 3D image files from the test print stage, we printed 10 ABS models on a Dimension Elite Printer (Stratasys, Eden Prairie, MN, USA) and seven plastic-wood hybrid models on a BigBox 3D Printer (Chalgrove, UK) (Figure 2C). We had intended to print equal numbers of each, however, the printer using the Woodfill kept getting jammed and starting over, and seven was all we could print in the timeframe we had available. The printer jamming was due in part to the print material and due to errors in the file geometry that were not adequately resolved during the editing stage. In total, it took about eight hours to print the 10 ABS lizards plus an additional four hours to dissolve the scaffolding. It took nearly five days to print the seven plastic-wood hybrid models (due to the printer jamming), and the scaffolding needed to be cut off by hand using an Exacto knife which took about an hour for all seven models. If the printer had not jammed, it would have taken two hours per model to print.

It was quite difficult to thread the narrow floral wire (26 gage, Panacea Products, Columbus, OH, USA) through the ventral holes in both Woodfill and ABS of models. The tube we made was curved, and in hindsight it should have been straight through the lizard midsection. Instead, we wrapped the wire around the midsection of the bodies with two long ends hanging off the ventral side. We then held the models by the wire and dipped them in melted plasticine clay (Craft Smart, Irving, TX, USA) to completely cover all parts of the body and the wire wrapped around the midsection. After the clay solidified (about 30 minutes), we folded the wire and wrapped each lizard in aluminum foil for transport to the field.

In total, our time investment from scanning to printing was relatively low: it took 20 hours from scanning the specimen to our first test print. Additional manipulations to the image took an additional 40 hours (an undergraduate working 5 hours/week for 2 months). Although we had to troubleshoot issues with our image and printing, the process was relatively easy due to the resources available at the 3D print centers (namely staff to mentor undergraduate on image software and troubleshoot printing issues), and that we did not need the surface to be an exact biological replica because we covered the models with clay.

### Field testing lizard models

Lizard models were deployed in developed (residential yard) and natural (national park) habitats on the island of Curaçao (Dutch Antilles) for 24-48h and then scored for predation. In both habitat types, models were anchored to tree branches, bushes, or rocks on the ground using the floral wire. We recorded evidence of predation from likely lizard and avian predators (Figure 2D). There were no differences in predation between ABS and Woodfill models (F_1, 30_=0.48, *P* = 0.49) or between small and large models (F_1, 30_=0.01, *P* = 0.93), yet more models were attacked in the natural versus developed habitat type (F_1, 30_=17.15, *P* < 0.001) (Figure 3). We attribute these differences in attack rates in habitat types to there being fewer resources available to predators in natural habitats during the height of the dry season compared to more abundant resources available in irrigated residential yards. Regardless, both material types were durable to the field conditions and none of our models experienced any structural problems during the experiment.

**Fig. 3.**
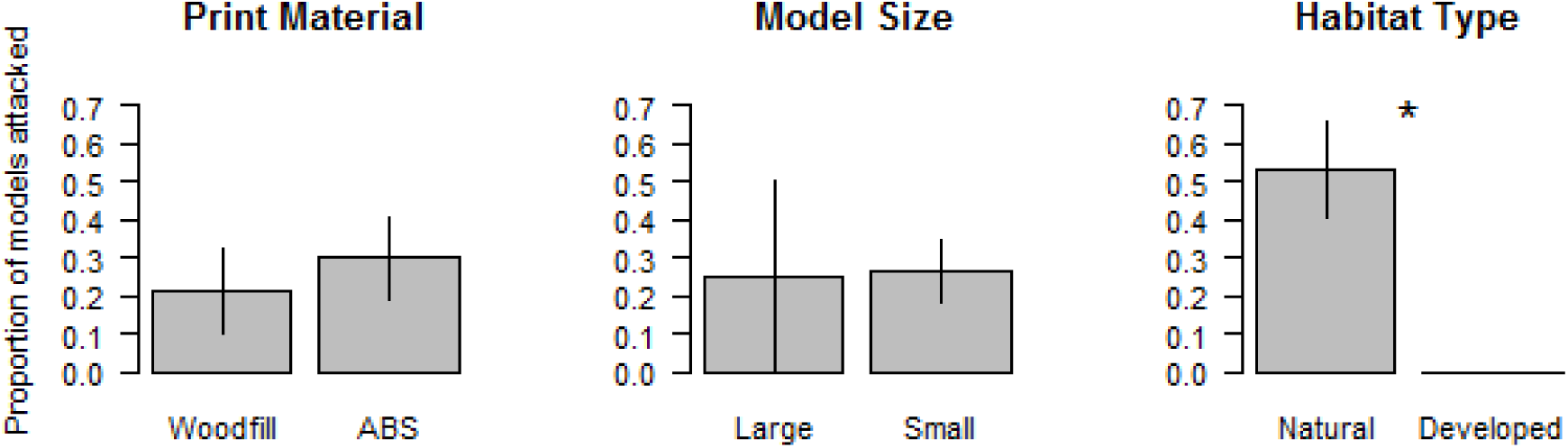
Proportion of clay-covered 3D printed models attacked during field predation trials. Bars are means +/- one standard error; * indicates significantly different means at *P*<0.01.

### Recommendations for using 3D printed models for field predation studies

Because the Woodfill models were cheaper and just as durable as the less sustainable ABS models, we would recommend using the Woodfill, or other similar plastics in comparable future studies, provided that the jamming issues we encountered during printing can be attributed to geometry errors in our file and not the Woodfill material itself. It should be noted that although we tested the models in extremely hot (>35°C) field conditions, we cannot comment on the durability of the two materials in rainy or very cold conditions. Initially, we believed the Woodfill would crumble more on the smaller model with narrower appendages, but this was not the case. Finally, our study took place over a 3-week period. It is possible that over longer time periods, the Woodfill would not be as durable as the ABS plastic.

## Discussion

### Reduce, Reuse, Recycle

While 3D printing can facilitate ecological research, the use of this technology must be weighed against its environmental and social costs. In general, 3D printing has the potential to reduce CO_2_ emissions and lead to more sustainable practices in the consumer manufacturing industry [53], yet there are still many less sustainable aspects to consider. Three-dimensional printing is much more energy intensive than 2D printing, and the most widely used materials in 3D printing, plastics, are fossil fuel based. Most 3D materials (e.g., plastic filament) are virgin and not derived from recycled post-consumer products, and if not disposed of and recycled properly, printed objects can exist in the environment for ages. The plastics used in printing can be toxic to aquatic organisms, especially resin-based printed objects [54]. The printing process itself generates extra waste due to printers jamming, misprinted models, and scaffolding necessary for more complex 3D objects. In addition, printing produces harmful emissions in the form of ultra-fine particles and volatile organic compounds [55, 56], which can be especially worrisome as many 3D printers are housed in indoor office settings [57]. With respect to the manufacturing of any plastic item, these negative aspects are not unique to 3D printing, they just become more obvious when one is directly involved in the manufacturing process. In our specific case, we chose 3D printed models for the speed at which they could be produced and their durability as we intend to use them in future experiments. Ecologists planning to incorporate 3D printing in research should strongly consider the negative impacts associated with 3D printing compared to the impacts of creating objects via other methods or not at all.

There are promising advances in the sustainability of 3D printing materials. Materials scientists are developing a range of filaments that are biodegradable, compostable, and made from recycled materials. For example, Eco-Filaments, such as WillowFlex (BioInspiration, Eberswalde, Germany), are made from plant-based resources and are completely compostable, even in residential compost bins. Other filament choices are made from recycled plastics like car dashboards, PET bottles, and potato chip bags (3D Brooklyn, Brooklyn, NY, USA; Refil, Rotterdam, the Netherlands). In fact, the cost of generating recycled plastic filament is often less than filament made from raw materials, prompting the establishment of a fair trade market for used plastic collected by waste pickers in the developing world (e.g., Protoprint Solutions, Prune, India) [58]. Non-plastic recycled filament options exists, such as filament made from the waste products of beer, coffee, and hemp production processes (3DFUEL, Fargo, ND, USA) as well as wood pulp [59]. Finally, because common print materials such as ABS plastic are not biodegradable or recyclable in municipal recycling centers, machines have been developed to recycle these plastics directly at the printing site [60]. These machines grind old prints and melt them into new filament that can be reused for printing (e.g. Filastruder, Snellville, GA, USA). Across sustainable options for print materials, we can attest to the durability of Woodfill for applications comparable to ours. For ecologists considering other sustainable print materials, most of these companies readily provide information about the durability of their products.

We stress that all 3D printing projects in ecological research should reduce, reuse, and recycle: *Reduce* the amount printed and the use of toxic print materials; *Reuse* printed objects and use materials made from post-consumer, waste materials; and *Recycle* printed objects by choosing materials that can be easily recycled, composted, or that are biodegradable. Planning a print job (Fig. 1) requires both careful estimation of the minimum number of replicates to print and smart design of geometry that minimizes or eliminates scaffolding, as scaffolding is usually discarded. Printing should be performed in well-ventilated environments where airborne toxins do not accumulate and harm personnel. The environmental toxicity of objects should be reduced by choosing materials with low toxic potential and reducing the toxicity of materials post-print. For example, exposure of resin-based printed objects to intense UV light can reduce their toxicity to aquatic organisms [54]. Printed objects should be reused in research as much as possible to avoid repeat printing, and print materials made from recycled material or materials that are recyclable or compostable should be used when possible. While most ecologists will not invest in their own 3D printing equipment and instead employ general-use academic (e.g., library) or commercial facilities, these environmental concerns can be communicated to the printing facilities so that they might adopt sustainable practices in their 3D printing for research.

## Conclusions

In conclusion, 3D printing technology has the promise to reduce the time and cost invested in creating custom materials used in ecological research, while at the same time increasing the ease at which collaborations occur within and outside the scientific community. Although there is a learning curve for developing 3D image files, there are ample online libraries of 3D files, plus tech savvy students and 3D printing center staff can be extremely helpful. Recent advances in print materials may reduce the footprint associated with this new technology. Overall, as with any new technology, ecologists must weigh the costs in terms of time and monetary investments into developing usable 3D materials for research versus other methods of generating those materials. If ecologists are in the position to commit the initial investment in securing printing resources and navigating the technological learning curve, the resulting ability to implement 3D printing into future studies could save time and money on the long term.

## Declarations

### Ethics approval

All work conducted involving live animals was in accordance with the Institutional Animal Care and Use Committee at Temple University (IACUC protocol #4614).

### Consent for Publication

Not applicable

### Availability of data and materials

The datasets used and/or analysed during the current study are available from the corresponding author on reasonable request.

### Author Contributions

JB and MH conceived of and designed the study; JB, BW, and MH developed the workflow; BW tested the workflow and designed the 3D model; JB and MH conducted the field experiment; JB, BW, STH, and MH wrote the manuscript and provided editorial advice. All authors read and approved the final manuscript.

## Acknowledgements

We thank J. Hample from the Digital Scholarship Center, S. Campbell from the Digital Fabrication Studio, and C. Denison from the Health Sciences Library Print Center all at Temple University for assistance with scanning and printing the lizard models. We thank M. Vermeij and S. Berendse from the Carmabi Foundation for logistical support in Curaçao.

## Additional Files

**Additional File 1.** Software for designing, modifying, and analyzing 3D files

**Additional File 2.** Online libraries of 3D imagery relevant for ecological research (as of 2017)

