## Supplementary Materials for "Benefits And Limitations Of Three-Dimensional Printing Technology For Ecological Research"

**Additional File 1 for:**

Table 1: Software for designing, modifying, and analyzing 3D files

| Name | Cost (in 2017) | Purpose | Edit 3D image | Generate image from photo | Build 3D image from scratch | Image analysis | Studies |
|---|---|---|---|---|---|---|---|
| 3D Lightyear | Free | Preparing 3D file to print; inputs STL/SLC files and prepares them for part building in other software | Yes | No | No | No | [1] |
| Agisoft PhotoScan | Reduced rate academic licenses | Photogrammetry software for processing digital images and generating 3D spatial data | Yes | Yes | No | Yes | [2] |
| Amira | Contact sales rep | Visualizing, manipulating, and understanding data from CT, MRI, microscopy, and | No | Yes | No | Yes | [3] |

| | | | | | | | |
|---|---|---|---|---|---|---|---|
| | | other imaging methods. | | | | | |
| AutoCAD (Autodesk) | Reduced rate academic licenses | Computer-aided design and drafting software for 3D modelling | Yes | No | No | No | [4] |
| Blender | Free | 3D image modeling, rigging, rendering | Yes | No | Yes | No | [5] |
| Checkpoint | Reduced rate academic licenses | 3D modeling, landmark collection and editing. | Yes | Yes | No | Yes | [6] |
| CTan | Free | 2D and 3D micro-CT dataset analysis and visualization | No | No | No | Yes | [7] |
| FreeCAD | Free | 3D modeling and modifying | Yes | No | Yes | No | [8] |
| Geomorph | Free | Geometric morphometric shape analysis in R environment | Yes | No | No | Yes | - |
| ImageJ | Free | Image enhancement, analysis, and editing | Yes | No | No | Yes | [5,7] |
| Inventor (Autodesk) | Free Academic License | 3D mechanical design, simulation, tooling | Yes | No | No | No | [5] |
| InVesalius | Free | Reconstruction of CT & MRI | No | No | No | Yes | [7] |
| Maya (Autodesk) | Free Academic License | 3D animation, modeling, rendering, simulation | Yes | No | Yes | No | This study |
| MeshLab | Free | Editing, cleaning, healing, inspecting, rendering and | Yes | No | No | No | [7] |

| | | | | | | | |
|---|---|---|---|---|---|---|---|
| | | converting unstructured 3D scans | | | | | |
| Meshmixer (Autodesk) | Free | 3D sculpting and image manipulation | Yes | No | Yes | Yes | [9] |
| MorphoJ | Free | Quantitative analysis of geometric morphometrics | No | No | No | Yes | [10,11] |
| OpenSCAD | Free | Focuses on CAD rather than artistic aspects of generating and manipulating 3D modelling | Yes | Yes | Yes | No | [12] |
| PhotoModeler Scanner | $2,495 | Alternative to 3D laser scanning to create 3D models from photographs | No | Yes | No | Yes | [10] |
| PhyloNimbus | Free | Landmarks & linear/curve measurement of 2D or 3D projects | No | No | No | Yes | - |
| Polyworks (Innovmetric) | Contact sales rep | Comprehensive 3D modeling software system with universal file formats | Yes | Yes | Yes | Yes | - |
| SketchUp (Google) | Reduced rate academic licenses | Designing 3D imagery from scratch | No | No | Yes | No | [13] |
| Solid Works | Contact sales rep | Most aspects of 3D image generation and manipulation | Yes | Yes | Yes | Yes | [12] |
| TinkerCad | Free | Highly accessible, web-based tool for designing new 3D models and, modifying existing 3D models | Yes | No | Yes | No | - |
