## Supplementary Materials for "Benefits And Limitations Of Three-Dimensional Printing Technology For Ecological Research"

**Additional File 2 for:**

Table 2: Online libraries of 3D imagery relevant for ecological research (as of 2017)

| Name | Website | Description |
|---|---|---|
| Cgtrader | https://www.cgtrader.com/3d-models | Repository of professional 3D models |
| GrabCAD | https://grabcad.com/ | Allows for printing straight from CAD; has a collaboration tool for working on group projects; also has library of free 3D imagery files |
| NIH 3D Print Exchange | https://3dprint.nih.gov/ | Provides scientifically accurate and medically applicable 3D models that are readily compatible with 3D printers. |
| Sketchfab | https://sketchfab.com/ | Platform for sharing 3D images on social media; also allows for 3D content to be embedded on webpages and social media |
| Thingiverse (by Makerbot) | http://www.thingiverse.com/ | Largest online community for sharing 3D models on an open platform |
| Dryad | http://datadryad.org/ | Repository for scientific files including 3D media |
| STL Finder | https://www.stlfinder.com/ | Search engine for 3D models |
